# Screening of Isolation and Culture Conditions of Lactic Acid Bacteria in Mongolian Horse Feces

**DOI:** 10.64898/2026.07.27.741107

**Authors:** Yanan Lin, Yang Zhang, Zhenqi Cao, Ying Wang, Jianfeng Cui, Ming Du, Jialong Cao, Xinlan Fang, Siqin Yun, Yajuan Weng, Shaofeng Su, Dongyi Bai, Yiping Zhao

## Abstract

**Objective:** This study aimed to establish an efficient isolation, culture, and preservation system for lactic acid bacteria from Mongolian horse feces by systematically evaluating the effects of sampling methods, culture conditions, media types, and cryopreservation protocols.

**Methods:** Fecal samples were collected using rectal and natural defecation methods, and bacteria were isolated by the spread plate technique, with isolates subcultured and identified via 16S rRNA gene sequencing. The performance of five commercial media (A–E), aerobic versus anaerobic conditions, and different glycerol concentrations (10% and 30%), temperatures (−20 °C and −80 °C), and storage durations (7, 15, and 30 days) were compared.

**Results:** Rectal sampling showed significantly higher lactic acid bacterial survival and greater species diversity (12 vs. 8). Anaerobic culture yielded substantially higher counts and diversity (707 strains, 12 species) compared to aerobic conditions (36 strains, 5 species). Medium C (M.R.S. AGAR) exhibited the best isolation performance, producing the highest number of strains (311) and species (11), with selective enrichment for both bacilli and cocci. Glycerol effectively improved strain survival during sample cryopreservation, with the optimal short-term storage (≤15 days) achieved at −20 °C with 30% glycerol. Functional screening of randomly selected strains from optimized conditions demonstrated promising growth, acid and bile salt tolerance, and antimicrobial activity.

**Conclusion:** Rectal sampling combined with anaerobic incubation using Medium C is recommended for optimal isolation. For samples that cannot be processed immediately, short-term preservation at −20 °C with 30% glycerol prior to isolation is effective. This study provides practical technical references for the acquisition and preservation of equine intestinal lactic acid bacteria resources.

## INTRODUCTION

Mongolian horses are recognized as one of the most ancient horse breeds and have contributed to the formation of many horse breeds worldwide. Currently, they are predominantly distributed in Mongolia and parts of China, including Inner Mongolia, Xinjiang, and the three northeastern provinces. Through long-term natural and artificial selection, Mongolian horses have acquired desirable traits such as strong adaptability, disease resistance, and tolerance to coarse feed (1). Zhao et al. (2) first characterized the gut microbiota of Mongolian horses by high-throughput sequencing and compared the fecal microbiota between Mongolian and thoroughbred horses, revealing significant differences in 5 phyla and 30 genera. The long-term exposure to cold, high-fiber grazing environments has shaped a distinctive gut microbial community in Mongolian horses, which plays a critical role in maintaining intestinal microecological balance, nutrient digestion and absorption, and immune regulation (3). Wen et al. (4) examined the gut microbiota of thoroughbred, Mongolian, and crossbred horses using second-generation high-throughput sequencing, and observed that the abundance of Bacillus was significantly higher in Mongolian horses than in the other two groups. Li et al. (5) compared the gut microbiota characteristics between Mongolian and Guizhou horses, and identified significant differences in 32 phyla and 718 genera between the two breeds, with the relative abundance of Lactobacillus and Lactococcus being significantly higher in Mongolian horses. Collectively, these sequencing-based findings point to a clear conclusion: Mongolian horses are not only important subjects for studying gut microbial diversity, but also represent a potential natural reservoir rich in lactic acid bacteria. Therefore, establishing an efficient isolation and culture system specifically for Mongolian horse-derived lactic acid bacteria is of considerable significance for the systematic exploitation of this unique microbial resource.

Among the various intestinal commensals, lactic acid bacteria represent an important group of probiotics. They are widely distributed in the animal gut, fermented foods, and natural environments, and have attracted considerable attention due to their multiple physiological functions, including modulation of host gut microbiota, enhancement of immune responses, and inhibition of pathogenic bacteria. Lactic acid bacteria are a general term for bacteria capable of metabolizing fermentable carbon sources and producing large amounts of lactic acid. They are typically Gram-positive cocci or bacilli (6).

Despite the considerable application potential of lactic acid bacteria in animal health, the food industry, and medical fields, current microbiological research faces a bottleneck—the “unculturability” of microorganisms. More than 70% of gut microbial species have not been cultivated, and the number of microorganisms that can be successfully cultured by humans remains limited(7). A review of the literature shows that large-scale cultivation of fecal samples has been conducted in mice (8), humans (9), and pigs (10), with corresponding gut microbial repositories having been established. However, extensive cultivation and characterization of gut microorganisms in equines have not been reported, with only a few studies on the isolation and culture of beneficial gut bacteria being available. For instance, Cao et al. (11) isolated a strain of Pediococcus pentosaceus M6 from equine sources, evaluated its probiotic properties in vitro, and demonstrated in a mouse model of DSS-induced colitis that the strain could alleviate weight loss, regulate inflammatory cytokine expression, modulate gut microbiota, and reduce intestinal barrier damage, suggesting its potential as a probiotic feed additive in animal husbandry.

Currently, there remains a lack of systematic technical protocols for the isolation and culture of lactic acid bacteria from horse fecal samples. This is reflected in several aspects: (1) whether the sampling method—such as rectal sampling versus natural defecation sampling—affects the isolation and culture of equine-derived lactic acid bacteria; (2) whether the post-sampling handling and preservation of fecal samples, including storage duration, temperature, and the need for protective agents, influence the isolation outcomes; and (3) insufficient evaluation of the applicability of commercially available selective media commonly used for lactic acid bacteria. These deficiencies may lead to low isolation efficiency and the omission of dominant strains. Therefore, the establishment of species-specific protocols for the isolation and culture of intestinal lactic acid bacteria in equines is urgently needed.

In vitro batch culture is a semi-representative model that provides a time-saving and cost-effective means of simulating physiological conditions (anaerobic, medium, 37°C) for culturing the gut microbiota (12). In this study, in vitro batch culture was employed to isolate, culture, and identify lactic acid bacteria from Mongolian horse feces. We systematically compared the effects of various procedural steps—from sampling to isolation, culture, and identification—on strain recovery, with the aim of determining optimal culture conditions for equine-derived lactic acid bacteria and establishing a protocol suitable for the isolation and culture of lactic acid bacteria from Mongolian horse feces. This study is expected to provide a theoretical basis and technical support for subsequent large-scale cultivation, fermentation process optimization, and the development of microecological preparations.

## MATERIALS AND METHODS

### 2.1. Sample Collection

Fresh fecal samples were collected from horses raised at the teaching farm of the Vocational and Technical College, Inner Mongolia Agricultural University, located in Tumed Right Banner, Baotou City, Inner Mongolia, China. Fresh fecal samples were collected from 20 healthy Mongolian horses (aged 3 to 8 years) using the rectal sampling method (Rectum) and the natural defecation sampling method (Feces). Each method was applied to 10 horses. The samples were placed into sterile centrifuge tubes, maintained at 4°C during transport, and subsequently processed under sterile conditions in a clean bench within the laboratory.

### 2.2. Reagents and Equipment

MRS broth, Lactobacillus Selective Agar, APT agar, MH agar, bovine bile salt, Bromocresol purple agar medium, and Gram staining kit were purchased from Qingdao Hope Bio-Technology Co., Ltd. (Qingdao, China). M.R.S. AGAR was obtained from Oxoid (Shanghai, China) via Shunyou (Shanghai) Biotechnology Co., Ltd. Lactobacillus agar medium was purchased from Guangdong Huankai Microbial Technology Co., Ltd. (Guangdong, China). Glycerol was obtained from Xilong Scientific Co., Ltd. (Shantou, China). Hydrochloric acid was purchased from Chengdu Jinshan Chemical Reagent Co., Ltd. (Chengdu, China). Bacterial genomic DNA extraction kit was obtained from Tiangen Biotech Co., Ltd. (Beijing, China). Premix Taq™ and DL2000 DNA Marker were purchased from TaKaRa (Dalian, China).

The anaerobic workstation (E500G) was manufactured by Gene Science (USA). The constant-temperature shaking incubator was obtained from Shanghai Yiheng Scientific Instrument Co., Ltd. (Shanghai, China). The upright microscope was from Nikon (Japan). The PCR thermal cycler was from BIO-RAD (USA). The gel imaging system was from Biorad (USA). The electrophoresis apparatus was from Beijing Liuyi Instrument Factory (Beijing, China). The automated growth curve analyzer (Bioscreen C°PRO) was from Oy Growth Curves Ab Ltd. (Turku, Finland). The pH meter (FE28) was from Mettler Toledo (Zurich, Switzerland).

### 2.3. Bacterial Strains

Staphylococcus aureus CMCC (B) 26003, Salmonella CMCC (B) 50071, and Escherichia coli ATCC 25922 were purchased from Beijing Zhongke Quality Inspection Biotechnology Co., Ltd. (Beijing, China).

### 2.4. Sample Processing

The fresh fecal samples obtained from 10 horses by rectal sampling (Rectum) and from another 10 horses by natural defecation sampling (Feces) were pooled separately in sterile bags and homogenized. One portion of each pooled sample was immediately subjected to the isolation and culture of lactic acid bacteria. The remaining portion was supplemented with glycerol at final concentrations of 0%, 10%, or 30% (v/v), and stored at either −20 °C or −80 °C. Isolation and culture of lactic acid bacteria from these frozen fecal samples were performed, followed by colony counting, on days 7, 15, and 30 of storage.

### 2.5. Isolation and Culture of Lactic Acid Bacteria

The samples were serially diluted ten-fold, and 100 μL aliquots of the 10⁻⁴, 10⁻⁵, and 10⁻⁶ dilutions were spread onto five different prepared media. The five media were Lactobacillus Selective Agar (Medium A) (13), APT agar (Medium B) (14), M.R.S. AGAR (Medium C) (15), Bromocresol purple agar (Medium D), and Lactobacillus agar (Medium E). All media were prepared strictly according to the manufacturers’ instructions. The inocula were evenly spread using a sterile spreader, with three replicates per group. The plates were then incubated under anaerobic (in an anaerobic workstation) and aerobic (in an incubator) conditions at 37 °C for 48 h. Based on the results obtained from the fresh fecal samples, the optimal culture conditions were selected and applied to the frozen fecal samples, using the same ten-fold serial dilution and spread plate method.

### 2.6. Subculture and Preservation of Lactic Acid Bacteria

After colony formation, record the colony data on the plate. Single colonies were picked based on their morphological characteristics and purified by streak plating. The well-grown single colonies were inoculated into MRS broth, and after three generations of subculture, the cells were centrifuged for collection, which was then used for the extraction of the bacterial DNA. The viable cell numbers of the primary (P), first (F), second (S), and third (T) passage strains were recorded. The survival rate was calculated using the following formula: survival rate (%) = (N₁ / N₀) × 100%, where N₁ represents the viable cell count after subculture and N₀ represents the viable cell count before subculture.

### 2.7. Morphological Identification of Lactic Acid Bacteria

Preliminary identification of the isolates was performed by Gram staining and microscopic observation of cellular morphology. The numbers of cocci and bacilli were recorded. Gram staining was carried out according to the manufacturer’s instructions provided with the kit (16).

### 2.8. Molecular Identification

Genomic DNA was extracted from the isolates using a bacterial genomic DNA extraction kit, and the 16S rRNA gene was amplified using a PCR thermal cycler. The universal bacterial primers 27F (5′-AGAGTTTGATCCTGGCTCA-3′) and 1492R (5′-GGTTACCTTGTTACGACTT-3′) were used for amplification. The PCR reaction mixture consisted of 0.8 µL of 27F, 0.8 µL of 1492R, 12.5 µL of Premix, 3 µL of template DNA, and 7.9 µL of ddH₂O, in a total volume of 25 µL. PCR reaction conditions: 95℃ for pre-denaturation for 5 minutes; 95℃ for denaturation for 30 seconds, 56℃ for annealing for 1 minute, 72℃ for extension for 1 minute and 30 seconds, for a total of 30 cycles; stored at 4℃.The amplified products were sent to Beijing Liuhe Huada Gene Technology Co., Ltd. (Beijing, China) for sequencing.

Upon obtaining the complete DNA sequences, BLAST homology searches were performed against the NCBI database (https://www.ncbi.nlm.nih.gov/) to determine the taxonomic affiliation of the isolates. The 16S rRNA gene sequences of the corresponding type strains were downloaded from the NCBI database. A phylogenetic tree was constructed using MEGA11 software with the Neighbor-Joining method (17).

### 2.9. Determination of Probiotic Properties of Lactic Acid Bacteria

Growth curve determination: Ten lactic acid bacteria strains were randomly selected from each of the following groups: rectal sampling, natural defecation sampling, aerobic culture, and anaerobic culture. Using the fully automatic growth curve analyzer, the sample was cultured at 37℃ for 24 hours. The absorbance at 600nm was measured every 2 hours to evaluate its growth performance.

Acid tolerance test: Using the fully automatic growth curve analyzer, the cultures were incubated at 37°C for 24 hours. The absorbance at 600nm was measured every 2 hours to evaluate the acid tolerance of the strains in media with pH values of 2.0 and 3.0.

Bile salt tolerance test: The strains were inoculated into MRS liquid medium with bile salt concentrations of 0.1% and 0.2%. The cultures were incubated at 37℃ for 4 hours. The viable bacterial count was determined using the dilution method and the plate plating technique. The survival rate was calculated using the following formula: survival rate (%) = (lg N₁ / lg N₀) × 100%, where lg N₁ is the logarithm of viable cell count after bile salt treatment and lg N₀ is the logarithm of viable cell count before treatment (18).

Antimicrobial activity test: Antimicrobial activity was assessed using the Oxford cup method. Selected lactic acid bacteria strains (10⁸ CFU/mL) were co-cultured with Staphylococcus aureus, Salmonella, and Escherichia colion MH agar plates. With three wells filled with lactic acid bacteria culture and one with sterile MRS broth as a negative control. Plates were incubated at 37 °C for 24–36 h, and inhibition zone diameters were measured to evaluate antimicrobial effects (19).

### 2.10. Statistical Analysis

All data are presented as mean ± standard deviation (SD). Statistical analyses were performed using SPSS software (version 23.0, IBM Corp., Armonk, NY, USA) with one-way ANOVA followed by Fisher’s least significant difference (LSD) test for multiple comparisons. Graphical representations were created using GraphPad Prism (version 9.0, GraphPad Software, San Diego, CA, USA).

## RESULTS

### 3.1. Effect of Sampling Method on the Isolation and Culture of Lactic Acid Bacteria

Fresh fecal samples collected by rectal sampling (Rectum) and natural defecation sampling (Feces) were subjected to spread plate isolation, yielding a total of 7,016 bacterial isolates. Of these, 3,451 primary isolates were obtained from Rectum and 3,565 from Feces. As shown in Figure 1, the survival counts of the first and third passage strains from rectal sampling were significantly higher than those from natural defecation sampling (p < 0.01). From the second passage onward, the survival of the strains tended to stabilize.

**Figure 1.**
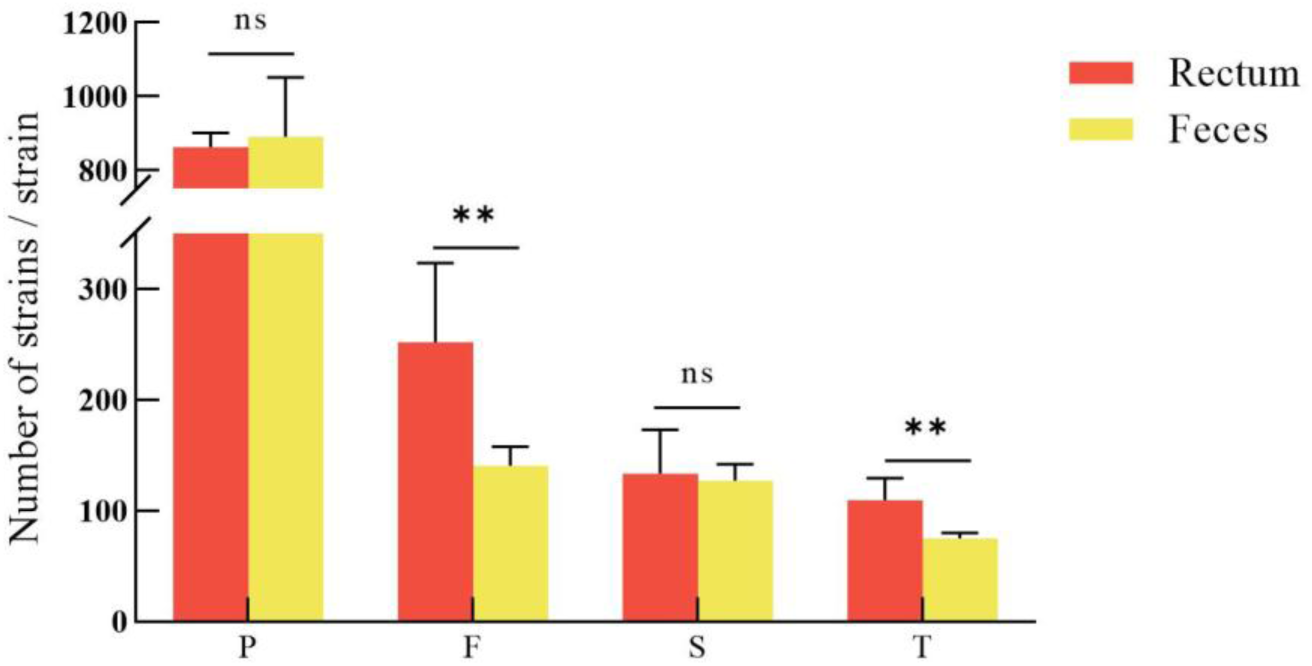
Comparison of the number of strains isolated and cultured by different sampling methods. ^*^ p<0.05, ^**^ p<0.01, ^ns^ p >0.05. P, Primary strain; F, First generation strain; S, Second generation strain; T, Three generations of strains. The same as below

Third-passage strains were subjected to sequencing, and BLAST analysis was performed against the NCBI database (Figure 2). A total of 743 lactic acid bacteria strains were identified, of which 421 were obtained by rectal sampling and 322 by natural defecation sampling. These strains belonged to 6 genera and 12 species. The six genera were Ligilactobacillus, Weissella, Streptococcus, Lactobacillus, Limosilactobacillus, and Pediococcus.

**Figure 2.**
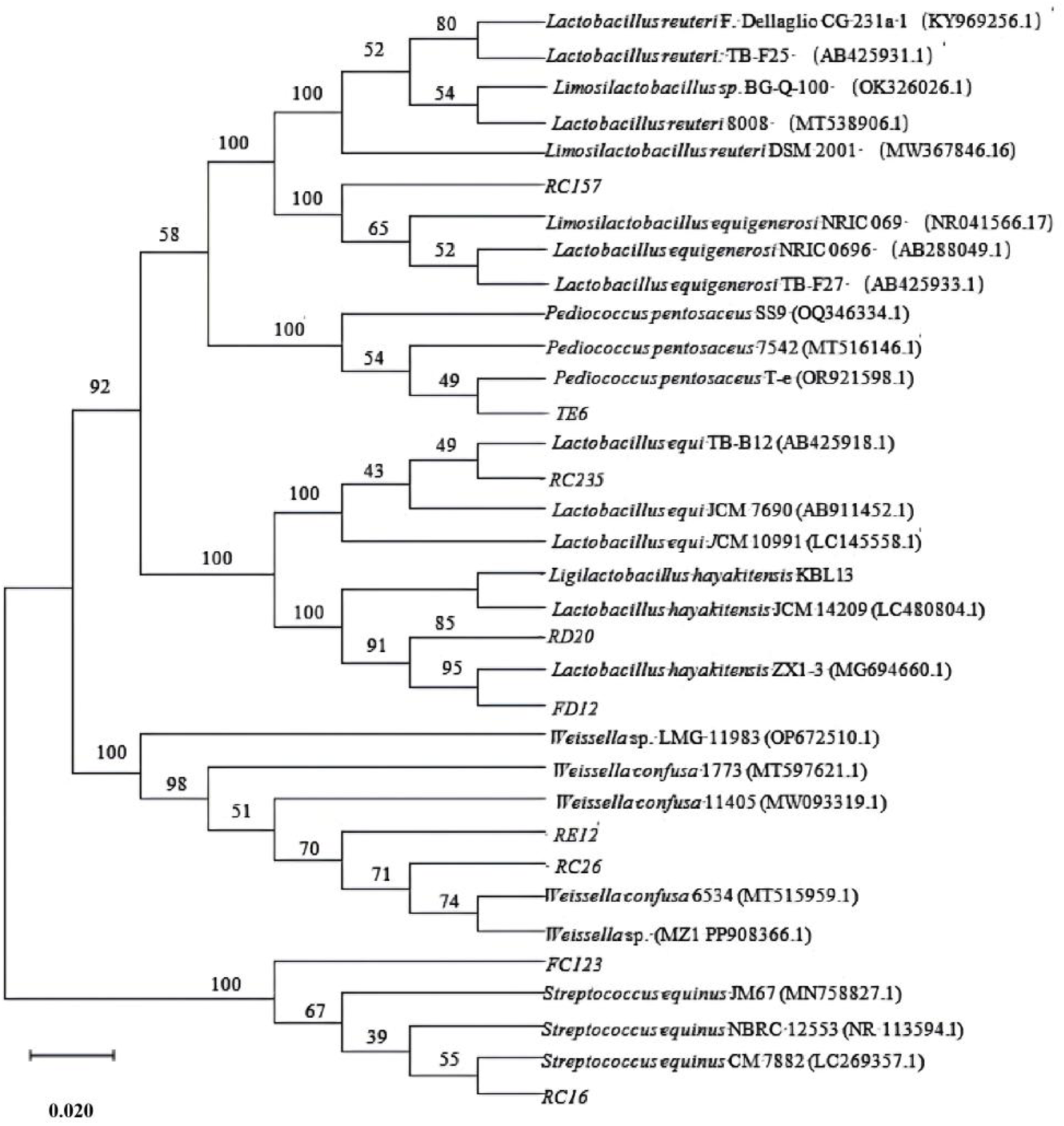
Phylogenetic tree of 16S rRNA gene sequence of lactic acid bacteria isolates.

As shown in Table 1, the Rectum method yielded a total of 12 types of lactic acid bacteria, while the Feces method yielded a total of 8 types of lactic acid bacteria. There was no significant difference between the two sampling methods in terms of the initial number of isolates obtained. However, rectal sampling resulted in significantly higher survival rates of the first and third passage strains (p < 0.01), and yielded a greater number and higher diversity of lactic acid bacteria species compared to natural defecation sampling.

**Table 1.** Comparison of lactic acid bacteria species and quantities isolated by Rectum and Feces methods.

| Species | Rectum | Feces |
| --- | --- | --- |
| *Ligilactobacillus equi* | 238 | 67 |
| *Weissella confusa* | 49 | 20 |
| *Weissella* sp. | 3 | 0 |
| *Streptococcus equinus* | 59 | 216 |
| *Streptococcus* sp. | 16 | 2 |
| *Streptococcus lutetiensis* | 1 | 3 |
| *Lactobacillus crispatus* | 26 | 0 |
| *Limosilactobacillus reuteri* | 20 | 0 |
| *Limosilactobacillus equigenerosi* | 1 | 4 |
| *Limosilactobacillus fermentum* | 2 | 0 |
| *Pediococcus pentosaceus* | 4 | 5 |
| *Ligilactobacillus hayakitensis* | 2 | 5 |

### 3.2. Effects of Aerobic and Anaerobic Conditions on the Isolation and Culture of Lactic Acid Bacteria

As shown in Figure 3, the number of strains isolated under anaerobic conditions was significantly higher than that under aerobic conditions (p < 0.01). Species identification revealed that only 36 strains belonging to 5 species were isolated under aerobic conditions. In contrast, 707 strains belonging to 12 species were isolated under anaerobic conditions (Table 2). These results indicate that both the number and species diversity of lactic acid bacteria isolated under anaerobic conditions were significantly higher than those under aerobic conditions (p < 0.01).

**Figure 3.**
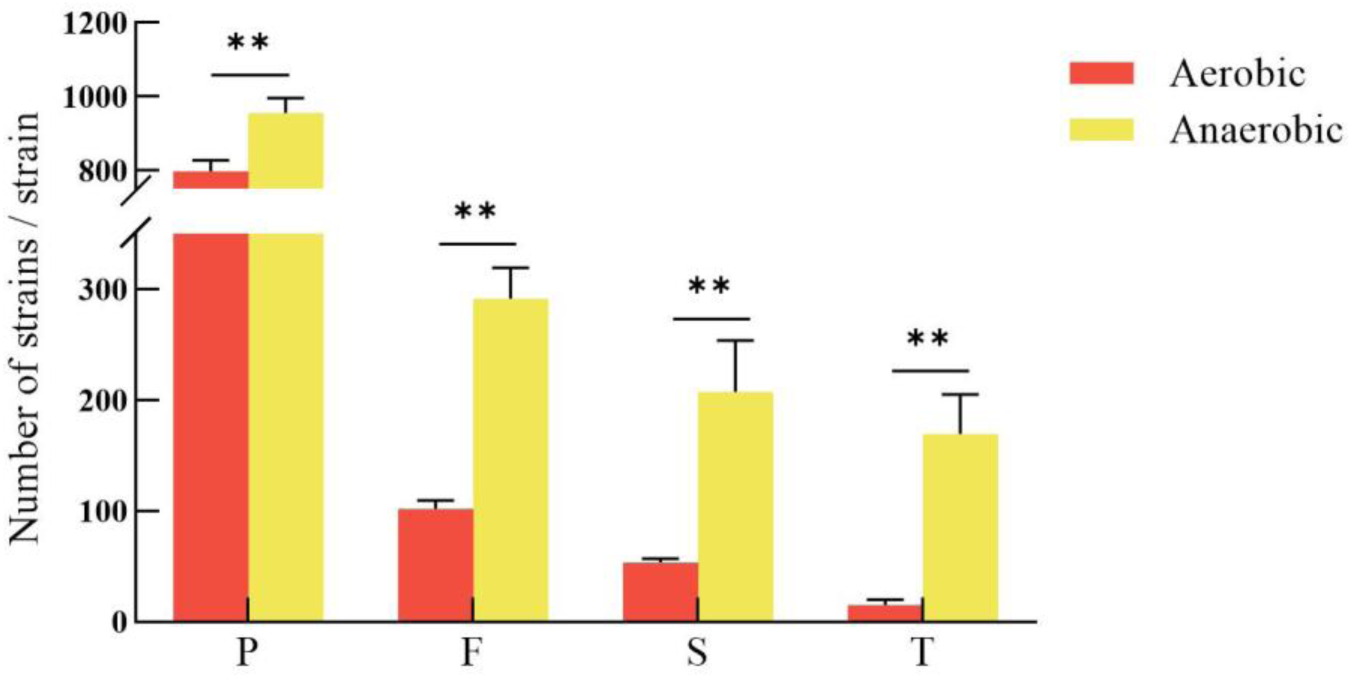
Comparison of the number of strains isolated and cultured under aerobic and anaerobic conditions. * p<0.05, ** p<0.01.

**Table 2.**
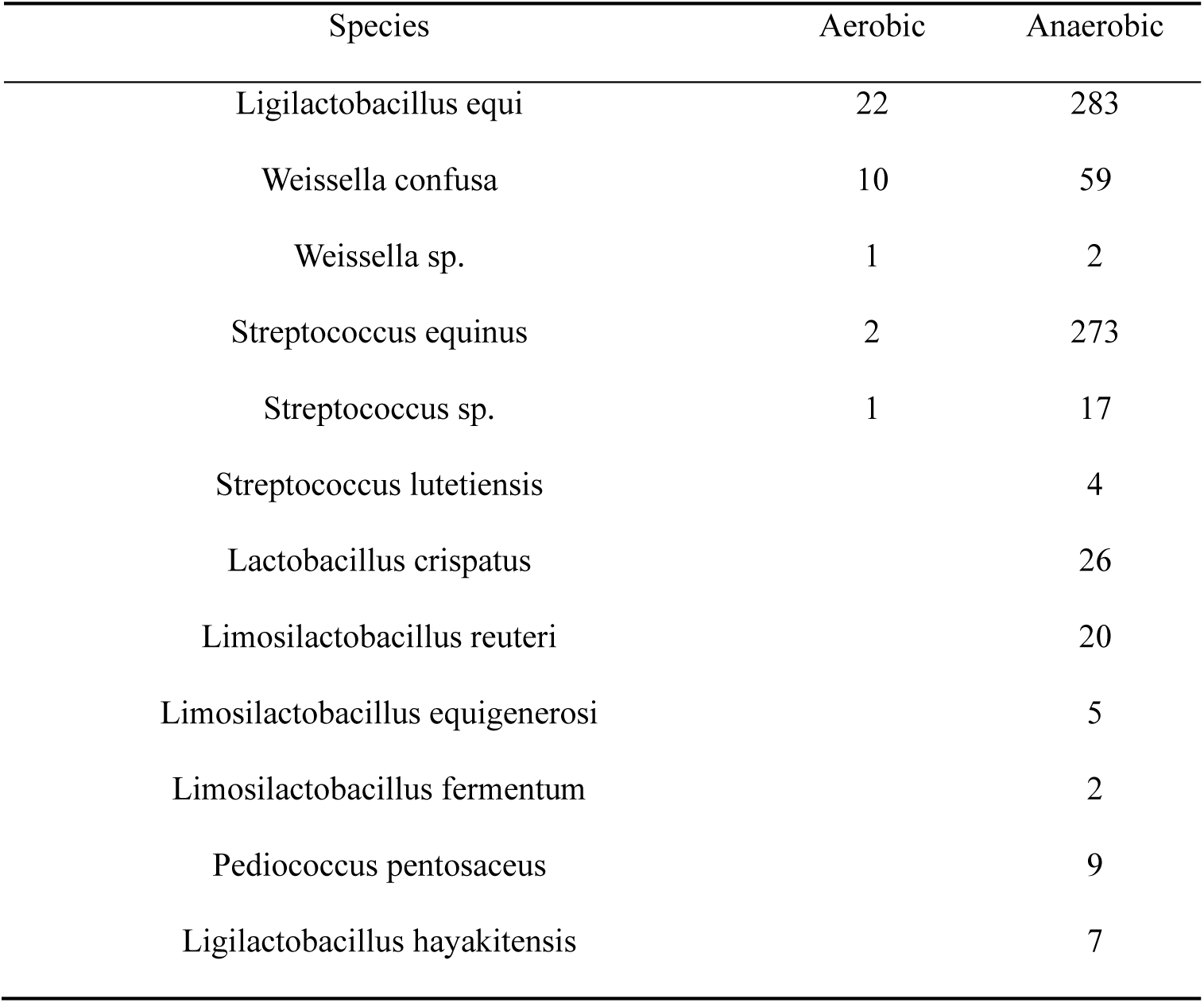
Comparison of lactic acid bacteria species and quantities isolated by Aerobic and Anaerobic.

### 3.3. Effects of Different Media on the Isolation and Culture of Lactic Acid Bacteria

As shown in Figure 4 and 6, Medium A yielded 122 strains belonging to 7 species; Medium B yielded 39 strains belonging to 3 species; Medium C yielded 311 strains belonging to 11 species; Medium D yielded 151 strains belonging to 7 species; and Medium E yielded 120 strains belonging to 9 species. The number of lactic acid bacteria obtained on Medium C was significantly higher than that on the other media (p < 0.01).

**Figure 4.**
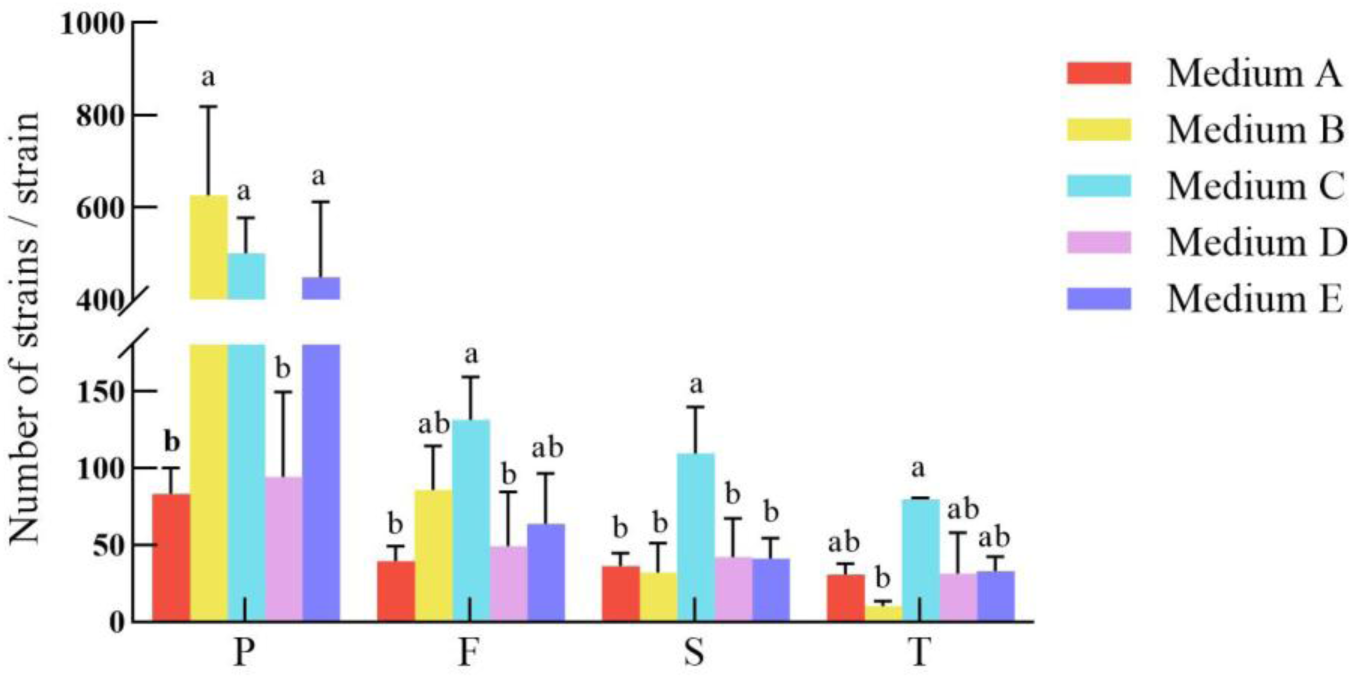
Comparison of the number of strains isolated and cultured in different media Vlues with different letter superscripts mean significant difference (p>0.05; While with the same or no letter superscripts mean no significant difference (p>0.05).The same as below.

**Figure 5.**
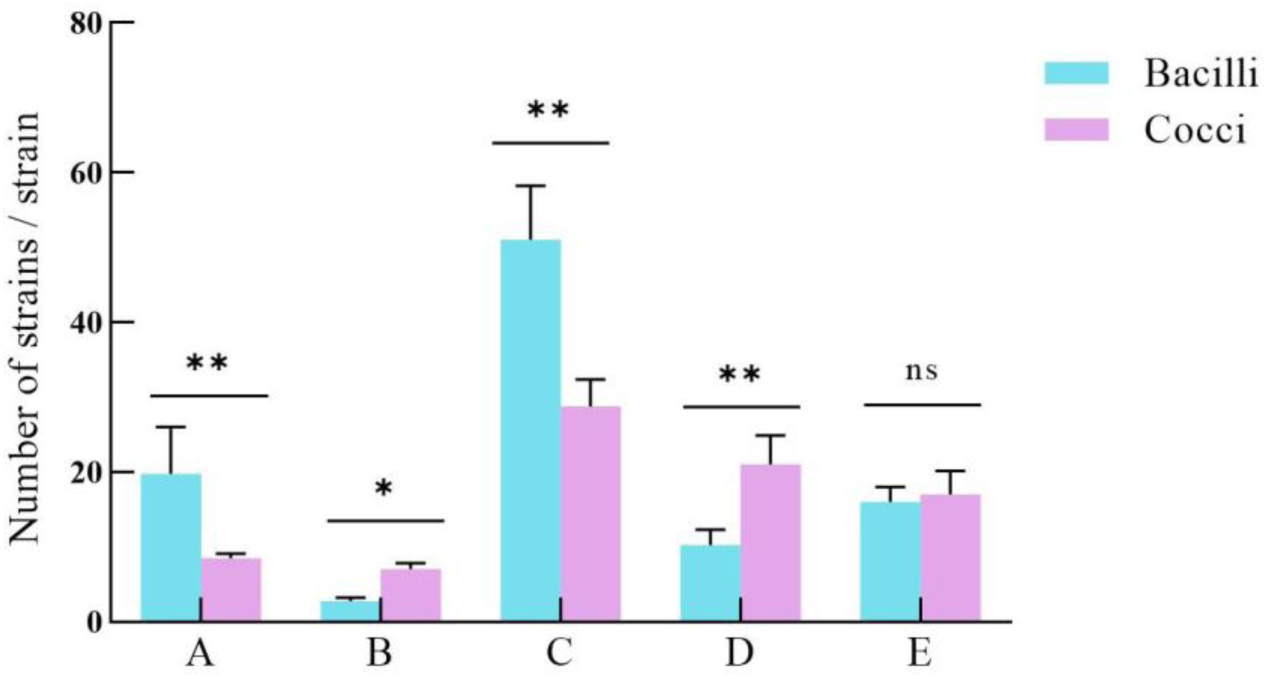
Comparison of the number of bacilli and cocci isolated and cultured in different media. * p<0.05, ** p<0.01, ^ns^ p>0.05.

**Figure 6.**
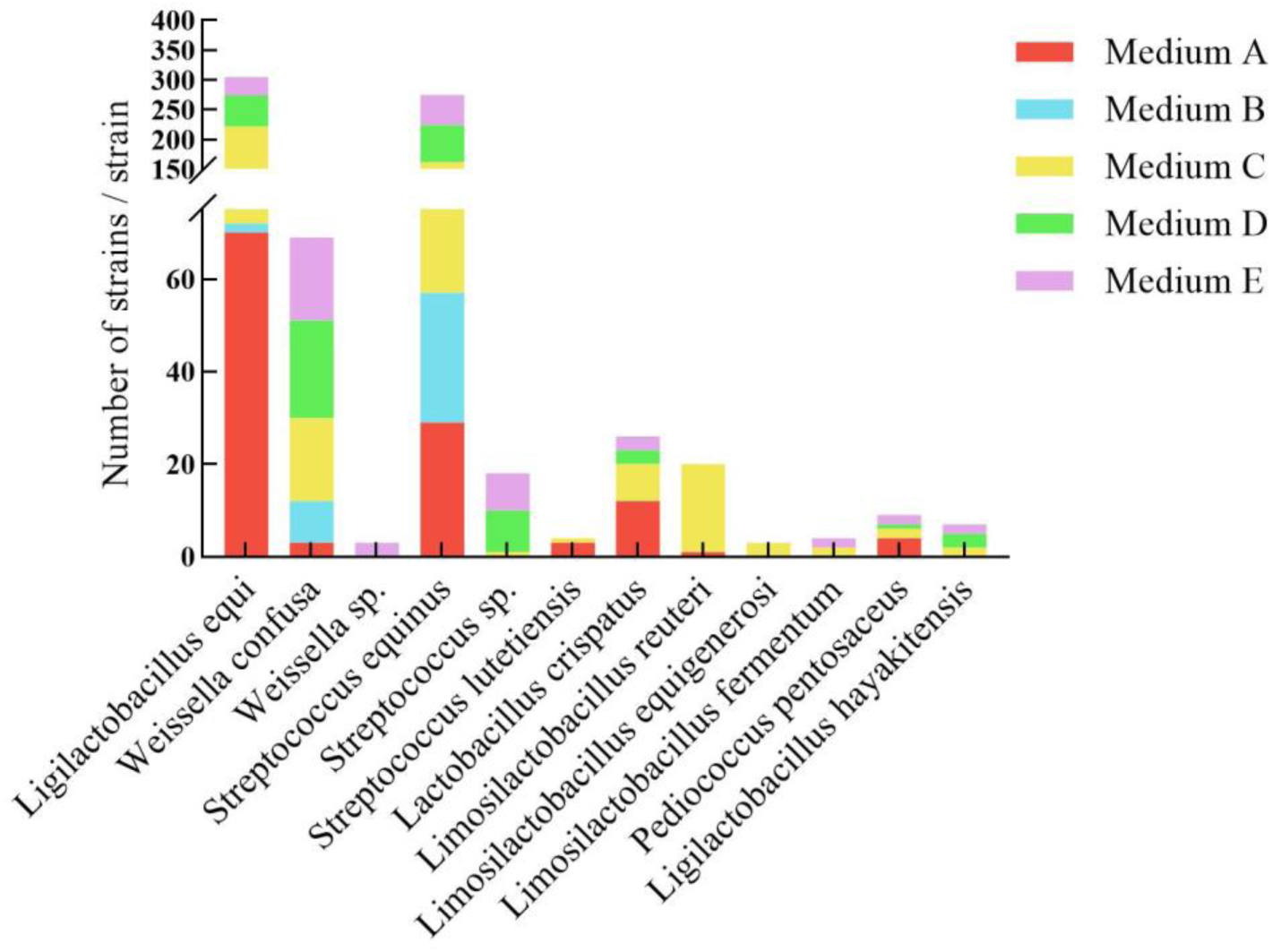
Comparison of the types and quantities of lactic acid bacteria isolated and cultured in different media.

The results showed that the different media exhibited selectivity toward bacilli and cocci (Figure 5). Media A and C favored the growth of bacilli, with the number of bacilli isolated being significantly higher than that of cocci (p < 0.01). In contrast, Media B and D favored cocci, with the number of cocci significantly exceeding that of bacilli (P < 0.01). On Medium E, the numbers of bacilli and cocci were similar, with no significant difference (p≥ 0.05).

### 3.4. Effect of Fecal Sample Cryopreservation on Subsequent Isolation and Culture of Lactic Acid Bacteria

The survival rate of culturable strains from fecal samples without glycerol protection declined sharply after freezing, whereas the addition of glycerol significantly improved survival. With increasing freezing duration, the number of culturable strains gradually decreased, with a maximum of 22 strains obtainable at day 30. The duration of freezing, storage temperature, and protective agent concentration all influenced the isolation outcomes, with short-term storage (≤ 15 days) at −20 °C with 30% glycerol yielding the best recovery of a large number of strains (Figure 7).

**Figure 7.**
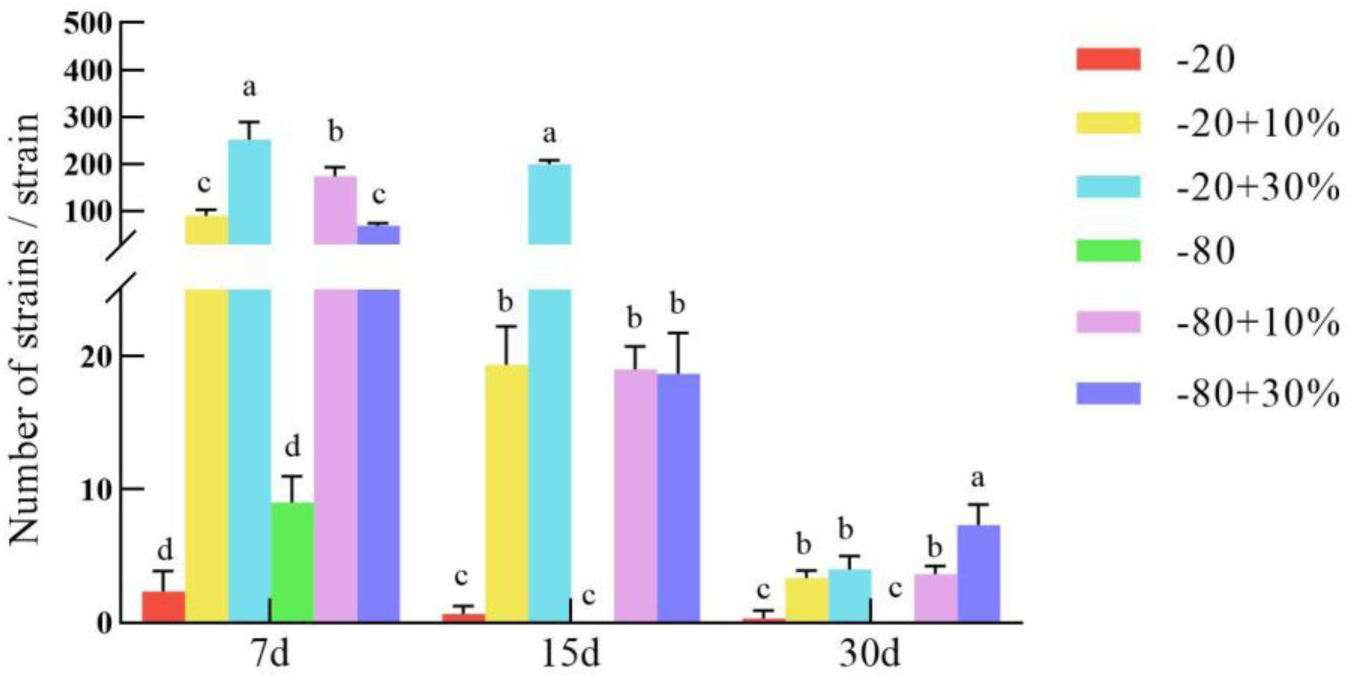
Comparison of the number of strains isolated and cultured after freezing. ^a-d^ Different letters indicate significant differences between mean values for a given behavior (p<0.05).

A two-way ANOVA was performed to evaluate the effects of storage temperature (−20 °C vs. −80 °C), glycerol concentration (0%, 10%, and 30%), and their interaction on the number of culturable strains after 7 and 15 days of cryopreservation (Table 3). At both time points, the main effect of temperature was not significant (p > 0.05), indicating that temperature alone did not determine strain recovery under short-term storage. In contrast, the main effect of glycerol concentration was highly significant (p < 0.001), confirming that the presence and level of cryoprotectant critically affected strain survival. Furthermore, the interaction between temperature and glycerol concentration was also highly significant (p < 0.001), suggesting that the optimal glycerol concentration depended on the storage temperature. Specifically, at −20 °C, 30% glycerol was more effective than 10%; at −80 °C, 10% glycerol was superior to 30%. At day 30, colony counts across all treatment groups dropped to very low levels (fewer than 10 colonies), precluding meaningful statistical analysis.

**Table 3.** Two-way ANOVA results for the effects of temperature, glycerol concentration, and their interaction on the number of culturable strains from fecal samples after 7 and 15 days of cryopreservation.

| Source of Variation | df | Day 7 | Day 15 |  |  |
| --- | --- | --- | --- | --- | --- |
|  |  | F-value | P-value | F-value | P-value |
| Temperature (T) | 1 | 2.84 | 0.103 |  | 0.088 |
| Glycerol concentration (C) | 2 | 68.53 | < 0.001 |  | < 0.001 |
| T × C interaction | 2 | 15.67 | < 0.001 |  | < 0.001 |

<sup>1)</sup> df, degrees of freedom. Significant effects are indicated in bold ( $p < 0.01$ ). The analysis was performed on data from day 7 and day 15; day 30 was excluded due to the absence of data for the 0% glycerol groups and overall low counts, which precluded valid interaction analysis.

### 3.5. Probiotic Performance Analysis of Lactic Acid Bacteria

As shown in Figure 8 during the first 0–4 h, most strains in the Rectum (R) and Anaerobic (AN) groups exhibited exponential growth, with OD₆₀₀ values exceeding 1.0. The stationary phase was reached at 6–8 h, with OD₆₀₀ values around 1.5. In terms of both growth rate and final biomass accumulation, the R group strains generally outperformed most of the Feces (F) group strains, and the AN group strains outperformed those in the Aerobic (AE) group.

**Figure 8.**
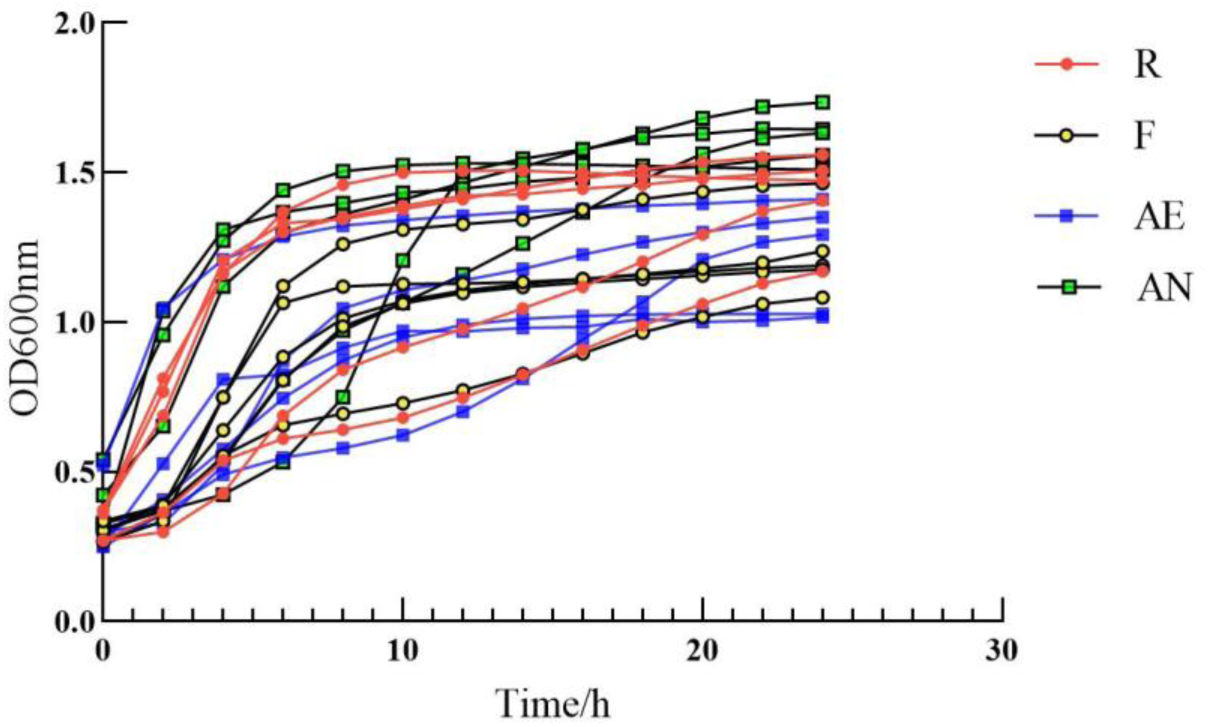
Partial growth curve of lactic acid bacteria.

The strains with the strongest growth performance from each group were selected for acid tolerance and bile salt tolerance tests. Strains RC138, RC157, and RC235 were able to grow under acidic conditions at pH 2.0 and pH 3.0 (Figure 9). As shown in Figure 10, the survival rates of RC138 and RC157 at bile salt concentrations of 0.1% and 0.2% were significantly higher than those of the F group strains (p < 0.05). All three strains (RC138, RC157, and RC235) were obtained by rectal sampling combined with Medium C. Among them, RC138 and RC157 were isolated under anaerobic conditions and identified as Ligilactobacillus equi and Limosilactobacillus reuteri, respectively, while RC235 was isolated under aerobic conditions and identified as Ligilactobacillus equi.

**Figure 9.**
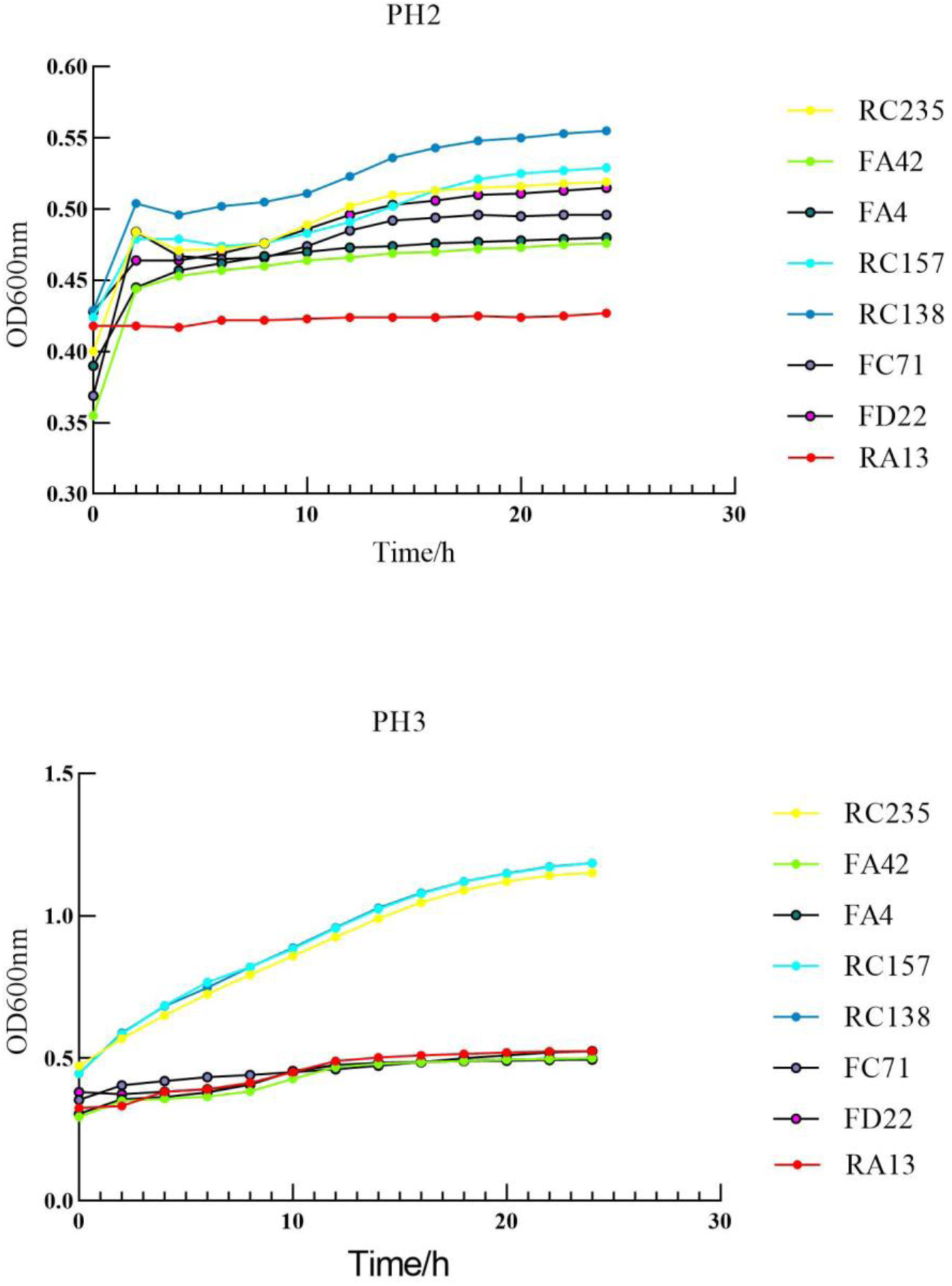
The growth curve of the strain in PH2 and PH3 acidic environment.

**Figure 10.**
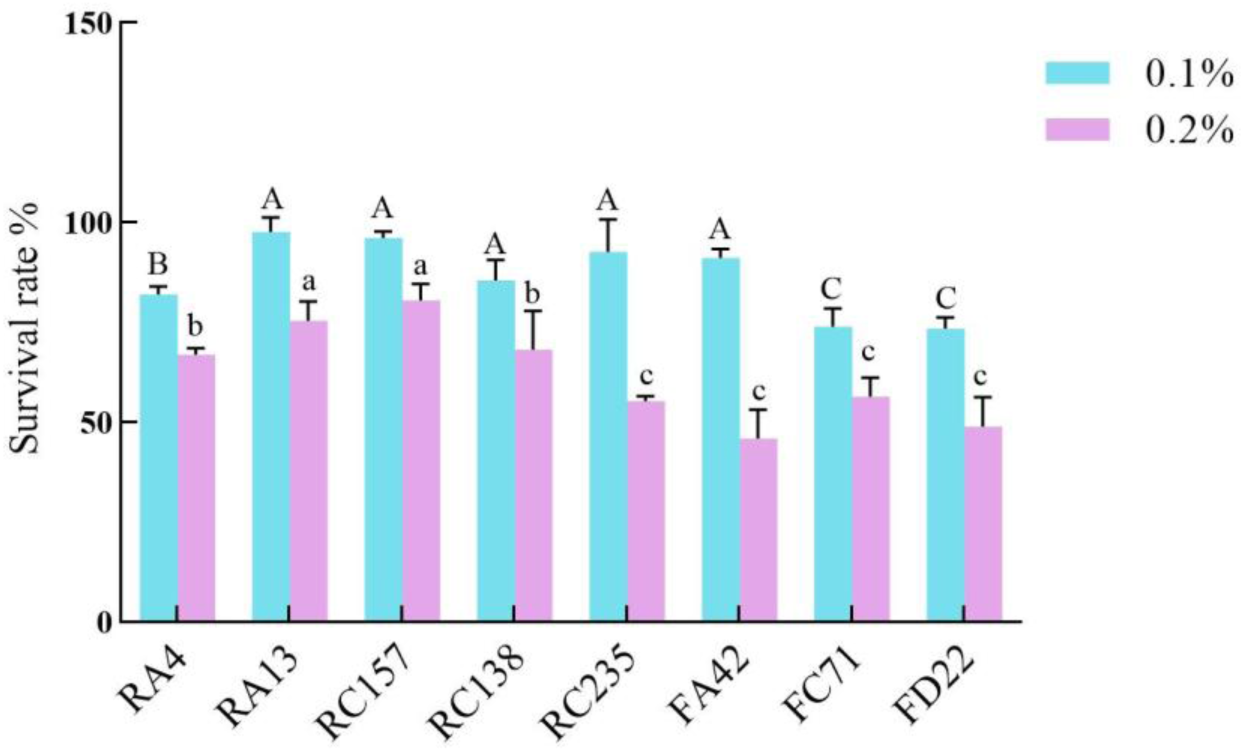
The survival rate of the strain after tolerance to 0.1 % and 0.2 % bile salt concentration for 4 h. ^a-c^ Different letters indicate significant differences between mean values for a given behavior (p<0.05).

The three strains with relatively strong tolerance, RC138, RC157, and RC235, were further evaluated for their antimicrobial activity against common pathogenic bacteria (Table 4). All three tested strains exhibited inhibitory activity against the selected pathogens to varying degrees, with RC157 showing relatively better activity against all three indicator strains.

**Table 4.** The antibacterial ability of the three strains against Staphylococcus aureus, Salmonella and Escherichia coli.

| Strain screening number | *Staphylococcus aureus* | *salmonella* | *Escherichia coli* |
| --- | --- | --- | --- |
| RC138 | 13.56±0.80b | 13.78±0.822b | 15.11±0.83a |
| RC157 | 15.64±1.41a | 17.29±0.52b | 15.59±1.39a |
| RC235 | 15.08±0.75a | 13.76±0.855a | 15.36±0.12a |

<sup>1)</sup> In the same column, values with different letter superscripts mean significant difference ( $p < 0.05$ ) ; While with the same or no letter superscripts mean no significant difference ( $p > 0.05$ ).

## DISCUSSION

Mongolian horses, as one of the renowned local breeds in China, harbor abundant microbial resources and represent an ideal equine breed for in-depth research on equine-derived lactic acid bacteria preparations. In recent years, several studies have applied diverse microbial isolation and culture methods combined with high-throughput identification techniques to isolate a large number of microorganisms from the intestines of humans, mice, and pigs, thereby enriching strain resources and providing a foundation for deciphering microbial “dark matter” as well as for subsequent functional and applied research on gut microbiota (20). To date, no reports have been found in the literature regarding optimized culture systems specifically for the isolation of lactic acid bacteria from the equine intestine.

The application of culturomics to isolate gut microorganisms from fecal samples enables the conversion of sequenced genes into living entities, which may lead to the discovery of novel species and unknown functions, and consequently elucidate strain-specific functions and mechanisms. The isolation and culture of lactic acid bacteria involve multiple steps, from sample collection to sequence data analysis, each of which is critical. Sampling, storage, and processing of specimens may all substantially influence the analytical outcomes. Ideally, specimen collection should be performed according to standardized protocols and should not introduce bias to the gut microbiota analysis results.

Regarding sampling methods, Santiago et al. (21) recommended homogenizing fecal samples during collection and transporting them to the laboratory for processing within 24 h (22). Radhakrishnan et al. (23) compared paired rectal swab and fecal samples from healthy individuals and found that the two sampling methods showed no significant differences in key alpha and beta diversity, major phyla abundance, and inferred gut functionality, with excellent correlation in functional prediction (Pearson’s r = 0.9217). Metabolomic analysis also revealed overall excellent correlation between the two methods. These findings are consistent with our results, in which no significant difference was observed between rectal sampling and natural defecation sampling in the initial number of isolates obtained. However, rectal sampling resulted in significantly higher strain survival rates during subculture and yielded a greater diversity of lactic acid bacteria species (12 species vs. 8 species). This may be attributed to the exposure of feces to oxygen, environmental microorganisms, and potential physical stress during natural defecation, which may inactivate or compromise the culturability of certain oxygen-sensitive or physiologically fragile lactic acid bacteria (24). Rectal sampling better preserves the original state of fecal microorganisms, particularly those that are strictly anaerobic or poorly oxygen-tolerant, which is crucial for obtaining a complete and authentic reservoir of intestinal lactic acid bacteria.

Nogacka et al.(25) isolated four lactobacilli strains (two Lactobacillus acidophilus and two Ligilactobacillus equi) from Asturcón horses and evaluated their effects on gut microbiota composition and metabolic activity using in vitro fecal batch fermentations. In the present study, a total of 743 lactic acid bacteria strains were isolated and purified using five media, belonging to 6 genera and 12 species. Among these, Ligilactobacillus equi was the predominant species in rectal samples (238 strains), whereas its proportion decreased substantially in naturally voided samples (67 strains). Ligilactobacillus equi is commonly found in the equine intestine and, as a dominant bacterial species, may alleviate Salmonella infection and regulate gut microbiota balance (26). In contrast, Streptococcus equinus was the dominant strain in naturally voided samples (216 strains). Streptococcus equinus is known to cause infectious diseases in horses, primarily manifesting as upper respiratory tract infections, particularly “strangles” (27). The marked differences in lactic acid bacteria species obtained by rectal sampling versus natural defecation sampling may be due to the differential tolerance of various lactic acid bacteria species to environmental stressors, including oxygen, ultraviolet radiation, temperature fluctuations, and contamination by environmental microorganisms (28). Streptococci generally exhibit stronger oxygen tolerance and thus are more likely to survive in naturally voided samples, whereas many Lactobacillus species are more sensitive and tend to be lost under exposure conditions. Natural defecation sampling may introduce selective bias, potentially leading to overestimation of the proportion of oxygen-tolerant strains and underestimation or even omission of important anaerobic lactic acid bacteria.

In this study, we also isolated 49 strains of Weissella confusa, 26 strains of Lactobacillus crispatus, 20 strains of Limosilactobacillus reuteri, and 9 strains of Pediococcus pentosaceus. Numerous studies have evaluated the probiotic properties of these species, demonstrating their important roles in food fermentation (29), human health (30), and disease prevention and treatment (31, 32). Comparative analysis of the probiotic properties of lactic acid bacteria obtained by rectal sampling versus natural defecation sampling revealed that strains obtained by rectal sampling exhibited superior performance in growth, acid and bile salt tolerance, and antimicrobial activity, further confirming that rectal sampling yields lactic acid bacteria strains of both greater quantity and better quality. These strains will be subjected to further in-depth investigation in our subsequent studies.

Secondly, the choice of culture conditions is a critical factor determining the success of isolation. Our results showed that the number and species diversity of lactic acid bacteria isolated under anaerobic conditions (707 strains, 12 species) were significantly higher than those obtained under aerobic conditions (36 strains, 5 species). This finding is highly consistent with the physiological characteristics of the intestinal environment, where most regions are anaerobic (33). Many lactic acid bacteria with potential probiotic functions, such as Lactobacillus crispatus and Limosilactobacillus reuteri, are strict or facultative anaerobes, and their growth may be inhibited or even terminated upon exposure to oxygen. Comparative analysis of the probiotic properties of strains obtained under aerobic versus anaerobic conditions revealed that strains isolated under anaerobic conditions exhibited superior performance in growth, acid and bile salt tolerance, and antimicrobial activity, further confirming that anaerobic culture yields strains of better quality. Therefore, the use of anaerobic culture techniques is an indispensable prerequisite for comprehensively exploring the diversity of intestinal lactic acid bacteria.

The selection of culture media exerts a pronounced “selective enrichment” effect on the isolation of lactic acid bacteria. In recent years, with the development of the market economy, the number of commercially available media brands has increased, and variations in raw materials and selectivity among different products may exist (23). Consequently, it is necessary to evaluate media to determine the most suitable formulations for the isolation of equine-derived lactic acid bacteria. Among the five media assessed in this study, Medium C (M.R.S. AGAR) performed best in terms of both the number and species diversity of lactic acid bacteria isolated. Comparative sampling of strains from the five media and screening for superior probiotic properties revealed that strains with better probiotic performance were all derived from Medium C. This may be attributed to the nutritional composition, pH, and redox potential of Medium C, which more broadly accommodate the growth requirements of various lactic acid bacteria in the equine intestine. It is noteworthy that different media exhibited distinct preferences for cellular morphology (bacilli vs. cocci). For instance, Media A and C favored the growth of bacilli, whereas Media B and D favored cocci. Such selectivity is closely associated with the specific nutritional requirements of different lactic acid bacteria species, including carbon sources, nitrogen sources, vitamins, and growth factors (33). Furthermore, certain species (e.g., a specific Weissella sp. that grew exclusively on Medium E) were isolated only on particular media, suggesting that the combined use of multiple media can more effectively avoid the omission of strains and thus provide a more comprehensive picture of the true lactic acid bacteria composition in samples.

It is worth noting that all strains obtained under different sampling methods and culture conditions exhibited varying degrees of loss during subculture, with the reduction rate from primary to first passage exceeding 70% in some cases. This may be due to the fact that the original strains are better adapted to their native environment, whereas laboratory media may not fully satisfy their growth requirements. In addition, prolonged laboratory cultivation may induce genetic changes in the strains, leading to reduced growth rates and metabolic capacity, which in turn affect the total bacterial count. Studies have shown that bacterial community diversity decreases significantly during continuous subculture, meaning that certain dominant strains may become predominant while others gradually disappear (34).

During bacterial subculture, cryopreservation is a core method for long-term strain preservation, but it may exert multiple effects on the biological characteristics of bacteria. During freezing, ice crystals formed from intracellular water can directly damage bacterial cells, leading to cell death and rupture, and consequently a reduction in cell numbers. Although glycerol is an effective cryoprotectant and was used in this study, it has certain limitations, including potential toxicity to some bacteria, with high concentrations potentially damaging cells. Other cryoprotectants, such as NADESs (natural deep eutectic solvents), may also reduce bacterial mortality during freezing and could be applicable for the cryopreservation of lactic acid bacteria (35). These issues warrant further investigation in subsequent studies. Numerous studies have demonstrated that sample preservation methods substantially influence the recovery of strains obtained by culture-dependent approaches. Long-term storage of samples is typically performed at −20 °C or −80 °C, which are regarded as the gold standards for sample preservation. For short-term storage (≤ 24 h), refrigeration at 4 °C is generally sufficient, whereas storage at room temperature is not recommended. Samples may also be stored after freeze-drying; however, this process can be damaging to cells and may greatly affect sample viability (36). The addition of protective agents, such as disaccharides and polyols, has been shown to improve the viability of bacteria in freeze-dried fecal samples (37). In our experiment on the effect of fecal sample cryopreservation on subsequent lactic acid bacteria isolation and culture, we further confirmed the necessity of glycerol as a cryoprotectant for maintaining the survival of lactic acid bacteria, and demonstrated that its protective effect is closely related to concentration, storage temperature, and duration. For short-term storage (7–15 days), samples supplemented with 30% glycerol and stored at −20 °C exhibited the best protective effect, still yielding a substantial number of culturable strains. This is mainly attributed to the ability of glycerol to penetrate cells, lower the freezing point, and reduce ice crystal formation, thereby mitigating mechanical damage to cell membranes and internal structures during freezing (38). Nevertheless, even under optimal conditions, the number and diversity of recoverable strains decreased substantially after 30 days of freezing, indicating that different lactic acid bacteria strains exhibit marked differences in their tolerance to freezing stress.

## CONCLUSIONS

This study systematically evaluated the combined effects of sampling methods, culture conditions, media types, and fecal sample cryopreservation protocols on the isolation and culture of lactic acid bacteria from Mongolian horse feces. The results demonstrated that the optimal strategy for isolating lactic acid bacteria from fresh fecal samples involved rectal sampling, anaerobic incubation, and the use of Medium C—which exhibited broad selectivity and high efficiency—supplemented with Medium A (targeting bacilli) and Media B and D (targeting cocci). In cases where immediate processing of samples is not feasible, glycerol can be used as a cryoprotectant for fecal samples, with short-term storage (≤ 15 days) at −20 °C with 30% glycerol yielding the best preservation outcome. This study provides detailed empirical support for standardizing the isolation and culture procedures of intestinal microorganisms, and establishes a solid foundation for subsequent probiotic screening and functional studies.

## CONFLICT OF INTEREST

The authors declare no conflicts of interest. The funders had no role in the design of the study; in the collection, analyses, or interpretation of data; in the writing of the manuscript; or in the decision to publish the results.

## AUTHORS’ CONTRIBUTION

Conceptualization, Y.L. (Yanan Lin) and Y.Z. (Yiping Zhao);

Methodology, Y.L.and S.S.(Shaofeng Su);

Validation, Y.L., Y.Z. (Yang Zhang), Z.C.(Zhenqi Cao),Y.W. (Ying Wang) and J.C. (Jianfeng Cui);

Formal analysis, Y.L.;

Investigation, Y.L., J.C. (Jialong Cao), X.F.(Xinlan Fang), S.Y.(Siqin Yun) and Y.W. (Yajuan Weng);

Resources, Y.L. and M.D.(Ming DU) data curation, Y.L., Z.C., Y.Z. Y.W. and J.C.;

Writing—original draft preparation, Y.L.;

Writing—review and editing, Y.Z. (Yiping Zhao) and D.B.(Dongyi Bao);

Visualization, Y.L.; supervision, Y.Z. and D.B.;

Project administration, Y.Z. and D.B.;

Funding acquisition, Y.Z. and D.B.

All authors have read and agreed to the published version of the manuscript.

## FUNDING

This research was funded by the Science and Technology Plan Project of Inner Mongolia Autonomous Region, grant number 2025KYPT0136, and the Fundamental Research Funds for the Directly Affiliated Universities of Inner Mongolia Autonomous Region, grant number BR261016.

## ACKNOWLEDGMENTS

The authors gratefully acknowledge the Equine Research Center for providing laboratory facilities and financial support. We also thank the teaching farm of the Vocational and Technical College, Inner Mongolia Agricultural University, for providing the fecal samples used in this study.

## SUPPLEMENTARY MATERIAL

Not applicable.

## DATA AVAILABILITY

The raw data supporting the conclusions of this article will be made available by the authors on request.

## DECLARATION OF GENERATIVE AI

During the preparation of this manuscript, the authors used ChatGPT (OpenAI, GPT-4o) for the purposes of language polishing and grammar checking. The authors have reviewed and edited the output and take full responsibility for the content of this publication.

## ETHICS APPROVAL

This study was conducted in compliance with the principles of the Declaration of Helsinki. The study protocol was reviewed and approved by the Institutional Animal Care and Use Committee of Inner Mongolia Agricultural University. All animal procedures were carried out in accordance with relevant institutional and national regulations.

## Notes

### Competing Interest Statement

The authors have declared no competing interest.

## REFERENCES

1. Zhang X, Liu Y, Li L, Ma W, Bai D, Dugarjaviin M. Physiological and Metabolic Responses of Mongolian Horses to a 20 km Endurance Exercise and Screening for New Oxidative-Imbalance Biomarkers. Animals : an open access journal from MDPI. 2025;15(9).

2. Zhao Y, Li B, Bai D, Huang J, Shiraigo W, Yang L, et al. Comparison of Fecal Microbiota of Mongolian and Thoroughbred Horses by High-throughput Sequencing of the V4 Region of the 16S rRNA Gene. Asian-Australasian journal of animal sciences. 2016;29(9):1345–52.

3. Lin Y, Qiri G, Du M, Dugarjaviin M, Cao J, Fang X, et al. Differential shaping of equine gut microbiota structure and function by breed and feeding regimen. 2026;Volume 17 - 2026.

4. Wen X, Luo S, Lv D, Jia C, Zhou X, Zhai Q, et al. Variations in the fecal microbiota and their functions of Thoroughbred, Mongolian, and Hybrid horses. Frontiers in veterinary science. 2022;9:920080.

5. Li Y, Lan Y. Characteristics and dynamic changes of gut microbiota in Mongolian horses and Guizhou horses. Frontiers in microbiology. 2025;16:1582821.

6. George F, Daniel C, Thomas M, Singer E, Guilbaud A, Tessier FJ, et al. Occurrence and Dynamism of Lactic Acid Bacteria in Distinct Ecological Niches: A Multifaceted Functional Health Perspective. Frontiers in microbiology. 2018;9:2899.

7. Almeida A, Nayfach S, Boland M, Strozzi F, Beracochea M, Shi ZJ, et al. A unified catalog of 204,938 reference genomes from the human gut microbiome. Nature biotechnology. 2021;39(1):105–14.

8. Liu C, Zhou N, Du MX, Sun YT, Wang K, Wang YJ, et al. The Mouse Gut Microbial Biobank expands the coverage of cultured bacteria. Nature communications. 2020;11(1):79.

9. Liu C, Du MX, Abuduaini R, Yu HY, Li DH, Wang YJ, et al. Enlightening the taxonomy darkness of human gut microbiomes with a cultured biobank. Microbiome. 2021;9(1):119.

10. Wylensek D, Hitch TCA, Riedel T, Afrizal A, Kumar N, Wortmann E, et al. A collection of bacterial isolates from the pig intestine reveals functional and taxonomic diversity. Nature communications. 2020;11(1):6389.

11. Cao J, Zhang J, Wu H, Lin Y, Fang X, Yun S, et al. Probiotic Potential of Pediococcus pentosaceus M6 Isolated from Equines and Its Alleviating Effect on DSS-Induced Colitis in Mice. Microorganisms. 2025;13(5).

12. Williams CF, Walton GE, Jiang L, Plummer S, Garaiova I, Gibson GR. Comparative analysis of intestinal tract models. Annual review of food science and technology. 2015;6:329–50.

13. Rogosa M, Mitchell JA, Wiseman RF. A selective medium for the isolation and enumeration of oral and fecal lactobacilli. Journal of bacteriology. 1951;62(1):132–3.

14. Evans JB, Niven CF, Jr. Nutrition of the heterofermentative Lactobacilli that cause greening of cured meat products. Journal of bacteriology. 1951;62(5):599–603.

15. De Man JC, Rogosa M, Sharpe ME. A MEDIUM FOR THE CULTIVATION OF LACTOBACILLI. Journal of Applied Bacteriology. 1960;23(1):130–5.

16. Beveridge TJ. Use of the Gram stain in microbiology. Biotechnic & Histochemistry. 2001;76(3):111–8.

17. Kumar S, Stecher G, Tamura K. MEGA7: Molecular Evolutionary Genetics Analysis Version 7.0 for Bigger Datasets. Molecular Biology and Evolution. 2016;33:1870–4.

18. Mañas P, Pagán R. Microbial inactivation by new technologies of food preservation. Journal of applied microbiology. 2005;98(6):1387–99.

19. Zhou Q, Xiao Q, Zhang Y, Wang X, Xiao Y, Shi D. Pig liver esterases PLE1 and PLE6: heterologous expression, hydrolysis of common antibiotics and pharmacological consequences. Scientific reports. 2019;9.

20. Xu MQ, Pan F, Peng LH, Yang YS. Advances in the isolation, cultivation, and identification of gut microbes. Military Medical Research. 2024;11(1):34.

21. Santiago A, Panda S, Mengels G, Martinez X, Azpiroz F, Dore J, et al. Processing faecal samples: a step forward for standards in microbial community analysis. BMC microbiology. 2014;14:112.

22. Gratton J, Phetcharaburanin J, Mullish B, Williams H, Thursz M, Nicholson J, et al. An optimized sample handling strategy for metabolic profiling of human feces. ACS Publications; 2016.

23. Radhakrishnan ST, Gallagher KI, Mullish BH, Serrano-Contreras JI, Alexander JL, Miguens Blanco J, et al. Rectal swabs as a viable alternative to faecal sampling for the analysis of gut microbiota functionality and composition. Scientific reports. 2023;13(1):493.

24. Lau JT, Whelan FJ, Herath I, Lee CH, Collins SM, Bercik P, et al. Capturing the diversity of the human gut microbiota through culture-enriched molecular profiling. Genome Medicine. 2016;8(1):72.

25. Nogacka AM, García A, G․ de los Reyes-Gavilán C, Arboleya S, Gueimonde M. In vitro assessment of horse-isolated strains of Lactobacillus acidophilus and Ligilactobacillus equi species for fecal microbiota modulation in horses. Journal of equine veterinary science. 2025;145:105341.

26. Zheng J, Wittouck S, Salvetti E, Franz C, Harris HMB, Mattarelli P, et al. A taxonomic note on the genus Lactobacillus: Description of 23 novel genera, emended description of the genus Lactobacillus Beijerinck 1901, and union of Lactobacillaceae and Leuconostocaceae. International journal of systematic and evolutionary microbiology. 2020;70(4):2782–858.

27. Taylor SD, Wilson WD. Streptococcus equi subsp. equi (Strangles) Infection. Clinical Techniques in Equine Practice. 2006;5(3):211–7.

28. Lagier J-C, Hugon P, Khelaifia S, Fournier P-E, la Scola B, Raoult D. The Rebirth of Culture in Microbiology through the Example of Culturomics To Study Human Gut Microbiota. Clinical microbiology reviews. 2015;28:237–64.

29. Park J-H, Ahn H-J, Kim S-g, Chung C-H. Dextran-like exopolysaccharide-producing Leuconostoc and Weissella from kimchi and its ingredients. Food Science and Biotechnology. 2013;22(4):1047–53.

30. Zalambani C, Rizzardi N, Marziali G, Foschi C, Morselli S, Djusse ME, et al. Role of D(-)-Lactic Acid in Prevention of Chlamydia trachomatis Infection in an In Vitro Model of HeLa Cells. Pathogens (Basel, Switzerland). 2023;12(7).

31. Smirnova E, Pimenov N, Ivannikova R. Immunotropic features of the sub-producer Limosilactobacillus reuteri. Veterinariya, Zootekhniya i Biotekhnologiya. 2023;3:103–10.

32. Ananda N, Suniarti D, Bachtiar E. The antimicrobial effect of Limosilactobacillus reuteri as probiotic on oral bacteria: A scoping review. F1000Research. 2023;12:1495.

33. Lagier J-C, Dubourg G, Million M, Cadoret F, Bilen M, Fenollar F, et al. Culturing the human microbiota and culturomics. Nature Reviews Microbiology. 2018;16(9):540–50.

34. Ricca DM, Ziemer CJ, Kerr BJ. Changes in bacterial communities from swine feces during continuous culture with starch. Anaerobe. 2010;16(5):516–21.

35. Jiang P, Li Q, Liu B, Liang W. Effect of cryoprotectant-induced intracellular ice formation and crystallinity on bacteria during cryopreservation. Cryobiology. 2023;113:104786.

36. Tan LL, Mahotra M, Chan SY, Loo SCJ. In situ alginate crosslinking during spray-drying of lactobacilli probiotics promotes gastrointestinal-targeted delivery. Carbohydrate Polymers. 2022;286:119279.

37. Tidjani Alou M, Naud S, Khelaifia S, Bonnet M, Lagier J-C, Raoult D. State of the Art in the Culture of the Human Microbiota: New Interests and Strategies. Clinical microbiology reviews. 2020;34.

38. Hirai M, Ajito S, Sugiyama M, Iwase H, Takata S-i, Shimizu N, et al. Direct Evidence for the Effect of Glycerol on Protein Hydration and Thermal Structural Transition. Biophysical Journal. 2018;115(2):313–27.

